# Fluctuations and the limit of predictability in protein evolution

**DOI:** 10.1101/2024.12.04.626874

**Authors:** Saverio Rossi, Leonardo Di Bari, Martin Weigt, Francesco Zamponi

## Abstract

Protein evolution involves mutations occurring across a wide range of time scales. In analogy with disordered systems in statistical physics, this dynamical heterogeneity suggests strong correlations between mutations happening at distinct sites and times. To quantify these correlations, we examine the role of various fluctuation sources in protein evolution, simulated using a data-driven energy landscape as a proxy for protein fitness. By applying spatio-temporal correlation functions developed in the context of disordered physical systems, we disentangle fluctuations originating from the initial condition, i.e. the ancestral sequence from which the evolutionary process originated, from those driven by stochastic mutations along independent evolutionary paths. Our analysis shows that, in diverse protein families, fluctuations from the ancestral sequence predominate at shorter time scales. This allows us to identify a time scale over which ancestral sequence information persists, enabling its reconstruction. We link this persistence to the strength of epistatic interactions: ancestral sequences with stronger epistatic signatures impact evolutionary trajectories over extended periods. At longer time scales, however, ancestral influence fades as epistatically constrained sites evolve collectively. To confirm this idea, we apply a standard ancestral sequence reconstruction algorithm and verify that the time-dependent recovery error is influenced by the properties of the ancestor itself. Overall, our results reveal that the properties of ancestral sequences—particularly their epistatic constraints—influence the initial evolutionary dynamics and the performance of standard ancestral sequence reconstruction algorithms.

## I. INTRODUCTION

Proteins play a key role in many biological processes that are essential for life. At the same time, they display a huge evolutionary flexibility, in that a large diversity of protein sequences can fold into the same structure and perform the same biological function. Such proteins are called ‘homologous’ and grouped into the same ‘protein family’ [1–4]. During evolution, a single ancestral protein belonging to a given family can diversify its amino acid sequence and, given enough time, explore a portion of the ‘neutral space’ of equivalently fit protein sequences [5]. Nevertheless, a random mutation in the amino acid sequence has a high probability of negatively impacting the structure and functionality of the protein [6–8]. Such kind of deleterious mutations are eliminated by natural selection, by which certain protein variants are favored over others due to their functional advantages: neutral (or even beneficial) mutations that maintain (or enhance) protein structure, stability, or function are more likely to persist in a population [9].

The picture gets more complex as one takes into account the concept of epistasis [10–18]: the effect of a mutation on the sequence fitness changes with the ‘background’ in which the mutation takes place, i.e. the amino acids that are present in the other sites of the protein. As a consequence, a mutation that would be deleterious in a particular background sequence can be instead beneficial in another (also called sign epistasis), allowing evolution to explore different pathways.

Understanding and characterizing the impact of epistasis in evolution requires careful experiments and modeling. In particular, recent developments have substantially increased the sequence divergence that can be reached by laboratory evolution experiments [19–23] A large amount of data is therefore now becoming available with more expected to come soon. Yet, the sequence diversity of natural evolution still remains out of reach of such experiments, thus leaving an unexplored gap in evolutionary time scales. In order to fill this gap, one can simulate the evolution of protein sequences *in silico* [24–29], relying on the data-driven approach that goes under the name of Direct Coupling Analysis (DCA) [15, 30]. DCA infers a fitness landscape (analogous to an energy function in the statistical physics vocabulary) starting from a Multiple Sequence Alignment (MSA) of natural homologs constituting a given protein family [31–35]. The energy function that results from this inference procedure can then be used to assign a probability to each sequence. The resulting landscape is explored by means of a biologically motivated Monte Carlo Markov-Chain (MCMC) algorithm, providing in silico evolutionary trajectories that quantitatively mimic experimental results [24–29, 36].

Within this framework, it has been recently shown [29] that different sites evolve with widely different time scales, which also depend on the background sequence, due to epistatic interactions. More precisely, sites that are more epistatically constrained need much longer times (or number of generations) to evolve (i.e. accumulate mutations) compared to less constrained sites [29]. Furthermore, whether a site is epistatically constrained or not depends on the rest of the sequence (the background) [37–39], which adds sequence-to-sequence heterogeneity on top of the site-to-site heterogeneity. This “heterogeneity of time scales” is strongly reminiscent of similar phenomena observed in disordered physical systems [40–50]. There, it has been shown that a large heterogeneity in time scales implies the presence of strong and highly collective space-time correlations between sites that slow down the dynamical evolution. Furthermore, it has been shown that the structural disorder in the initial configuration partially encodes future correlations, and can be used (possibly by machine learning tools) to predict the future dynamics [51–54], see [55] for a recent review. These are the main observations that motivated this work.

In this paper, we measure space-time correlations to explore how the heterogeneity of evolutionary time scales relates to the epistatic interactions within the ancestral sequence. We follow the dynamics of the Hamming distance, i.e., the number of accepted mutations with respect to the ancestral sequence. We characterize how this quantity fluctuates (i) between different evolutionary trajectories originating from the same ancestor, and (ii) between different ancestors. We observe a strong dependence of the dynamics on the ancestral sequence for short enough times, over which the first source of fluctuations is found to be subdominant with respect to the second. On the other hand, at long times the stochasticity of evolutionary trajectories takes over, and any memory of the ancestor is lost. This has important consequences, as it allows us to properly quantify the time scale over which it should be possible to reconstruct the ancestral sequence of a certain set of evolutionary trajectories, and how this time scale depends on the epistatic interactions in the ancestor itself. In fact, we find that the amount of epistatically constrained sites in the ancestral sequence determines this time scale: more epistatic sequences leave their trace on evolutionary dynamics for longer times. We then measure the correlations between the evolution of all pairs of sites at the time scale at which such correlations are stronger, finding a different pattern for each ancestral sequence, which is then reflected in the evolutionary dynamics. Finally, we show that the magnitude of epistatic interactions, which controls the stochasticity of the evolutionary trajectories, is also related to the linear response of the evolutionary dynamics to a change in selective pressure (controlled by the temperature in our Monte Carlo simulations). Hence, we show and quantify how more epistatically constrained ancestors lead to a more complex evolutionary dynamics over longer time scales, which is also more sensitive to perturbations of the environment.

## II. EPISTASIS IN BIOLOGICAL DATA

Epistasis plays a central role in shaping protein evolution, influencing both the structure of fitness landscapes and the fate of evolutionary trajectories, and as such it has been the subject of theoretical, computational and experimental studies, see Refs. [10–18] for a few recent reviews. It has been shown that some epistatic effects can be accounted for by a global non-linearity of the phenotype-fitness relation [56–59], but epistasis is also due to a network of more specific pairwise and higherorder interactions. While some of these interactions are sparse [18, 60–66], multiple studies suggest that the most relevant effects tend to emerge collectively, from the accumulation of numerous weak interactions between a single site and multiple other residues across the protein sequence [14, 15, 32, 38, 39, 67–70]. This form of distributed epistasis suggests that the evolutionary constraints on a given mutation are shaped by the broader sequence context, rather than by a few dominant interactions, highlighting the complexity of protein fitness landscapes.

Experiments have provided valuable insights into how mutations interact, but each methodology comes with inherent limitations. While some studies focus on measuring epistasis in a restricted mutational space, others provide broader datasets but lack information on evolutionary dynamics. Here, we summarize the main categories of experimental data available and discuss their relevance to our theoretical framework.

- Combinatorial mutagenesis [18, 63, 65, 71–74] experiments involving a small number of residues have demonstrated epistatic effects. However, these studies typically do not allow the accumulation of sufficient mutational effects to observe the largescale evolutionary patterns we investigate in this work.
- Deep mutational scanning (DMS) experiments across homologous wild-type proteins [22, 39, 75, 76], reveal epistatic interactions and can be used to explore the concept of site variability within a small region of the landscape. However, these experiments lack direct observations of evolutionary dynamics over multiple generations.
- Experimental studies tracking evolutionary dynamics in vitro are available [19–21, 23], but only a few cases, such as TEM-1 [19] and PSE-1 [20] *β*-lactamases, involve multiple wild-type homologs belonging to the same protein family, whose evolutionary dynamics can be directly compared. Moreover, these experiments do not extend far enough in sequence space to capture the long-term epistatic effects central to our study.

Given these limitations, we anticipate that future advancements in experimental techniques will provide richer datasets capable of testing the predictions outlined in our study. In the meantime, our theoretical framework serves as a guide for interpreting existing data and shaping expectations for future experimental work.

## III. METHODS

### A. Modeling evolution in silico

In order to mimic the evolution of protein sequences, we use the DCA model energy as a proxy for the fitness landscape. We start by building an MSA of naturally occurring sequences for each protein family we want to study. From the MSA we infer the parameters (fields and couplings) of the DCA model via Boltzmann Machine learning (bmDCA) [77, 78]. The resulting model assigns a probability *P* (*A*) = exp[−ℋ (*A*)]*/Z* to each sequence *A* = (*a*_1_, …, *a*_*L*_), with *L* the common length of aligned sequences in the MSA and *a*_*i*_ being a symbol that takes 21 possible values corresponding to the 20 natural amino acids plus the gap symbol needed for alignment. According to the statistical physics language, low energy

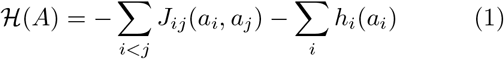

corresponds to high probability, hence high fitness. In this expression, epistatic interactions between different amino acids are represented via the pairwise couplings *J*_*ij*_(*a*_*i*_, *a*_*j*_) that have been found to be crucial in data-driven statistical models of biological sequences, cf. [15, 79]. Because training of these models has become a standard and well-documented procedure [77, 78], we do not give further details here.

Following [24, 25, 29], we consider a fixed initial sequence 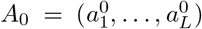 as the ancestor, and we let many trajectories evolve from it in parallel, hence restricting ourselves to a star phylogeny describing an ensemble of independent evolutionary trajectories of common initialization. Our goal is to characterize the statistical properties of the specific ancestral sequence and the sequence space accessible from it in a given evolutionary time. The sequence evolution is simulated by Monte Carlo dynamics. In this work, we use the Metropolis algorithm acting on amino acids for simplicity, and we verified that more refined sampling strategies taking into account amino-acid accessibility via the genetic code, insertions and deletions [24, 25, 29] produce qualitatively the same results, see the Supplemental Material (SM). As it was previously shown [79], at large times the generated sequences accurately reproduces many statistical features of the natural sequences used for the training, and have the same probability of being biologically functional in a given experimental platform; hence, the model is generative. Here, we are concerned with what happens at short and intermediate times, where the influence of the ancestral sequence is still important.

The results we report mainly concern the DNA-binding domain (DBD) protein family already studied in a similar setting in Ref. [29] and experimentally in Ref. [22], but for some results we also generalize to other protein families, in particular the WW domain (WW), Chorismate Mutase (CM), Aminoglycoside 6-N-acetyltransferase (AAC6), Dihydrofolate reductase (DHFR), Beta Lactamase (BL), and Serine Protease (SP) families. These families have been chosen because they span several chain lengths, and experimental data obtained either from Deep Mutational Scanning or by *in vitro* evolution are available. The procedures to construct the natural MSAs for these families are given in the SM.

### B. Measures of epistasis

By using the MSA of a given protein family as input data, and inferring the fitness landscape parameters via the DCA model, the authors of [29, 38] have been able to roughly classify the sites of any specific protein sequence belonging to the family in three categories: conserved, mutable, and epistatically constrained. In order to do so, they defined two site-mutability metrics.

- The Context-Independent Entropy (CIE) is obtained by computing the empirical frequency *f*_*i*_(*a*) of occurrence of amino acid *a* on site *i*, for every *a* and *i*, in the MSA obtained from the natural sequences. This is then used to compute a standard Shannon entropy as

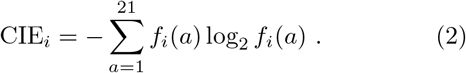

This quantity measures to what extent a site is variable or conserved across the input MSA.
- The Context-Dependent Entropy (CDE) is defined for site *i* in sequence *A* as

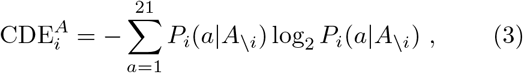

using the conditional probability *P*_*i*_(*a*|*A*_\*i*_) = *P* (*a*_*i*_ = *a*|*A*_\*i*_) of having amino acid *a* on site *i* given the rest of the sequence *A*_\*i*_ = (*a*_1_, …, *a*_*i−*1_, *a*_*i*+1_, …, *a*_*L*_). This quantity cannot be extracted directly from the input MSA and needs to be obtained from the model parameters; due to the epistatic couplings in the energy ℋ (*A*) it actually differs from the CIE. As indicated explicitly, the 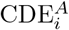 depends on the site *i* and the sequence *A*, hence on the context in which site *i* finds itself. As a matter of fact, this metric quantifies the local mutability of a site, i.e. its mutability within a certain reference-sequence context *A*_\*i*_.

A large value of CIE or CDE means that many mutations are allowed, while a small value means that only one or a few amino acids are tolerated.

In terms of these quantities, a site *i* in a given background sequence *A* can thus be classified as follows:

- *Mutable sites* have a large CDE and a large CIE. These sites can tolerate many mutations, both in the natural MSA and in the considered background.
- *Conserved sites* have a small CDE and a small CIE. These sites do not tolerate mutations, neither in the natural MSA nor in the considered background.
- *Epistatically constrained sites* have a small CDE and a large CIE. These sites display a large variability in the natural alignment, but only because the rest of the sequence is mutating at the same time. In fact, in the considered background, only one or a few amino acids are tolerated.

Note that it is very rare that a site has a CDE larger than the CIE, because generally speaking, fixing the background reduces the number of mutations that can be tolerated [38]. We also stress that this classification depends on the background *A*, and Ref. [29] has shown that epistatically constrained sites in a background can be mutable in another background and viceversa (while conserved sites tend to remain so in all backgrounds).

Furthermore, and most importantly for the present work, Ref. [29] has considered a given background *A*_0_ as the ancestral sequence, and starting from it has performed many parallel evolutions in silico, looking at how each site diversifies in the library of mutants obtained after a certain evolutionary time. It was found that mutable sites evolve very rapidly and quickly reach their asymptotic mutability, i.e. the CIE. Conserved sites remain so at any time during evolution, hence do not display interesting dynamics. Epistatically constrained sites, instead, are conserved at short evolutionary times, due to the epistatic constraints in the background of the ancestor, which cause a small CDE. But, as soon as the background sequence mutates enough, they can tolerate more mutations, asymptotically reaching the large CIE that they display in the natural alignment. Their mutation, however, is contingent on several mutations happening in the rest of the sequence, which often require a rather long evolutionary time to take place [29].

Along with the classification of sites discussed above, we also introduce a global measure of the level of epistasis in a given sequence *A*. Because each sequence has a different set of variable and epistatically constrained sites, we measure its overall level of epistatic constraints by averaging the CDE over sites,

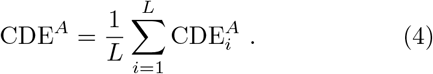

Hence, a more epistatically constrained sequence will display a lower CDE^*A*^.

### C. Dynamical fluctuations

Building on the previous work described above, and inspired by works on disordered systems [40–49], we devised a protocol (sketched in Fig. 1) to quantify how the amount of epistasis in the ancestral sequence impacts the subsequent evolution. We consider a series of ancestral sequences, taken from the natural MSA, let us call them *A*_0_, *B*_0_, *C*_0_, etc. Each ancestor has a different level of epistatic constraints, as measured by the average CDE introduced in Eq. (4), as illustrated in Fig. 1a.

**Figure 1.**
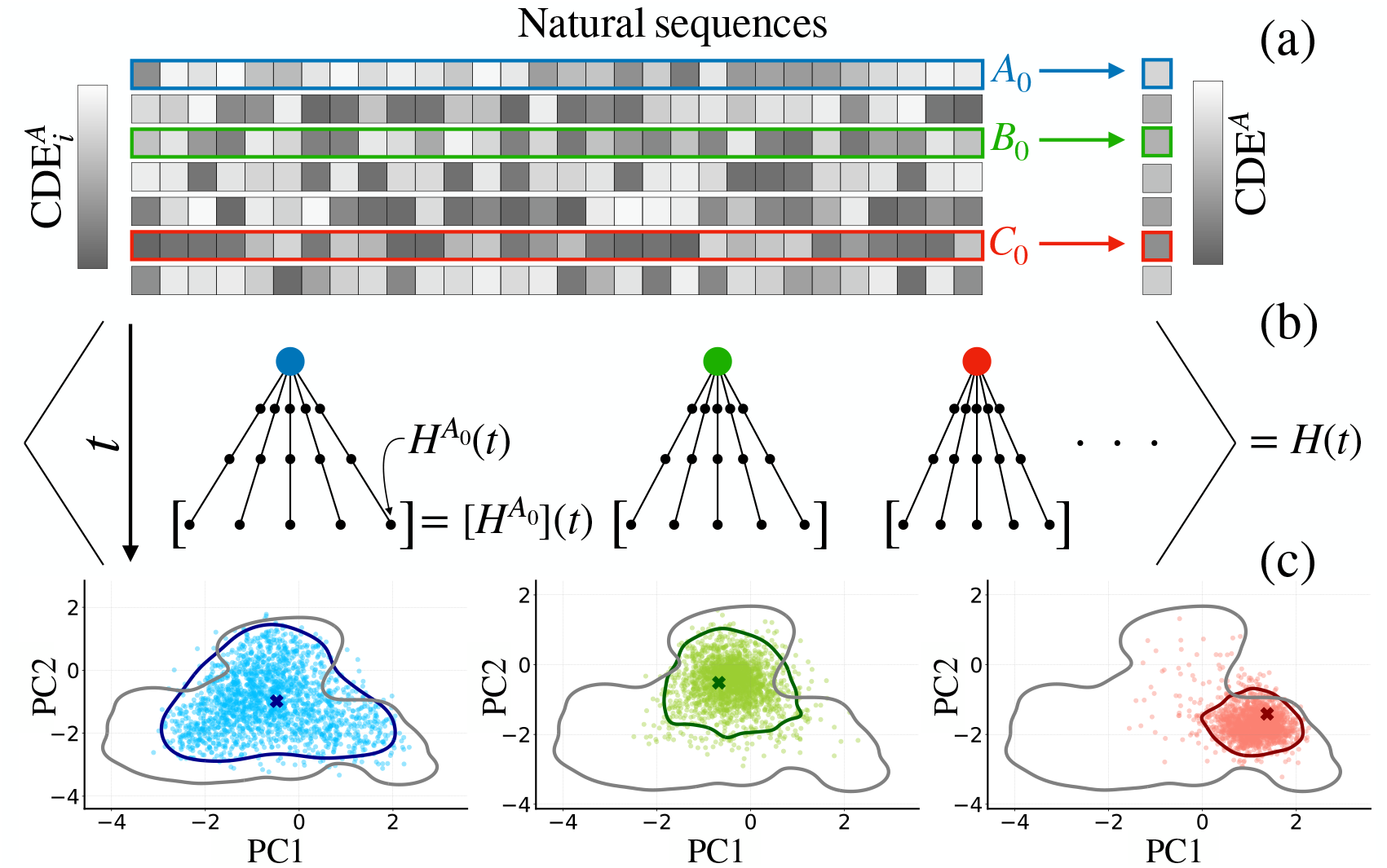
Schematic representation of (a) an MSA in which each site is colored based on its mutability as measured by the 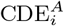 and (b) the protocol of our evolutionary dynamics with the relevant averages. (c) Depending on the epistatic constraints acting on the ancestral sequence and for equal evolutionary times, we observe a different level of diversity of the evolved sequences (colored), corresponding to a different exploration of the functional sequence space (grey).

Next, we consider a star phylogeny of independent MCMC evolutionary trajectories, all starting from the same ancestor and evolving in parallel (Fig. 1b). Hence, for each ancestor, we construct, as a function of evolutionary time, an MSA of descendants that mimic those obtained through in vitro evolution experiments. As a measure of diversity, we focus on the evolution of the Hamming distance, between the ancestor *A*_0_ and the MSA of evolving sequences at time *t*, 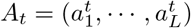, defined as

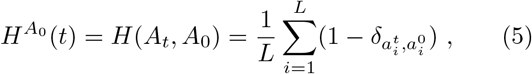

introducing the Kronecker delta symbol 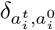, which is zero if site *i* is mutated between the two sequences *A*_*t*_ and *A*_0_, and one otherwise. This quantity corresponds to the ‘overlap’ in the disordered systems literature and to the number of accepted mutations in evolution. Other measures could be considered as well, but we focus on this one for simplicity. For a given ancestor, simulating many parallel evolutionary trajectories, we thus obtain a set of realizations of the random variable 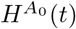.

As illustrated through a projection in PCA space in Fig. 1c, we observe that the level of epistasis in the ancestor determines the size of the portion of the fitness landscape explored by evolution. For comparable evolutionary time, ancestors with more epistatically constrained sites (e.g., sequence *C*_0_ in Fig. 1) lead to a less diverse set of evolved sequences. Conversely, ancestors with less epistatically constrained sites (e.g., *A*_0_ in Fig. 1) lead to a widely diverse set of evolved sequences.

To make this observation quantitative, we characterize the statistical properties of the resulting MSAs, and how they depend on the ancestor, by introducing two distinct averages and corresponding fluctuations, inspired from the disordered systems literature [45, 46, 48–50] and illustrated in Fig. 1b. Recall that the number of accepted mutations 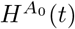 at evolutionary time *t* for fixed ancestor *A*_0_ is a random variabile, whose realizations depend on the stochasticity of the evolutionary trajectory.

First, we consider averaging over many evolutionary trajectories for fixed ancestor. We denote this average as [· · ·]. The variance of 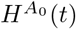 an then be defined as

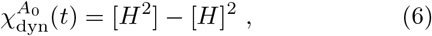

where the dependence of 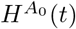 on *A*_0_ and *t* is omitted to simplify the notation. The quantity 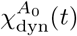 depends on the ancestor *A*_0_ and on time *t*.

- Second, we consider the average over different ancestors, which we define as ⟨· · · ⟩. In particular, we consider the average number of mutations 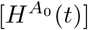 for a given ancestor, that measures the average diversity of the evolved MSA, and we measure how this quantity fluctuates from ancestor to ancestor via the variance

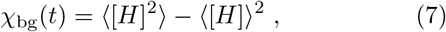

where again the dependence of 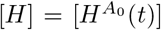 on *A*_0_ and *t* is omitted to simplify the notation. This quantity is the variance of [*H*] associated to the fluctuations in the background of the ancestral sequence *A*_0_, hence the suffix ‘bg’.

Note that the total variance of 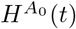 over both sources of randomness, i.e. the random choice of ancestor and the stochasticity of mutations along the evolutionary trajectory, can be decomposed as

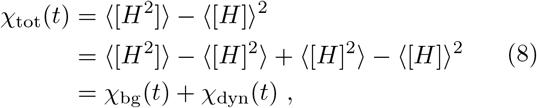

where

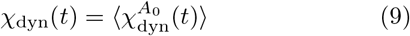

is the average of the ancestor-dependent 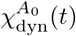 over *A*_0_. Using these quantities, which are called ‘dynamical susceptibilities’ in the physics of disordered systems, we can thus quantify the relative importance of the ancestor and the evolutionary noise in determining the diversity of the resulting MSAs at a fixed evolutionary time. In the SM we also study some alternative definitions of susceptibilities, based for example on the Hamming distance between two chains starting from the same ancestor. However, because our MCMC algorithm is time-reversible, these two quantities are related.

## IV. RESULTS

### A. Mutational dynamics and its fluctuations

The dynamical evolution of the Hamming distance 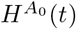 (number of accepted mutations with respect to the ancestor) is displayed as a function of the number of Monte Carlo sweeps in Fig. 2, with a sweep corresponding, on average, to one attempted mutation per site. More specifically, Fig. 2a shows the average 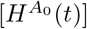 over a star phylogeny of 10^3^ parallel evolutions, for 200 choices of the ancestor *A*_0_ taken at random (with weights, see SM) from the DBD protein family. Fig. 2b reports the variance 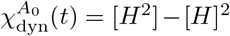 defined in Eq. (6) over the same phylogeny, and for the same initial sequences as in Fig. 2a.

**Figure 2.**
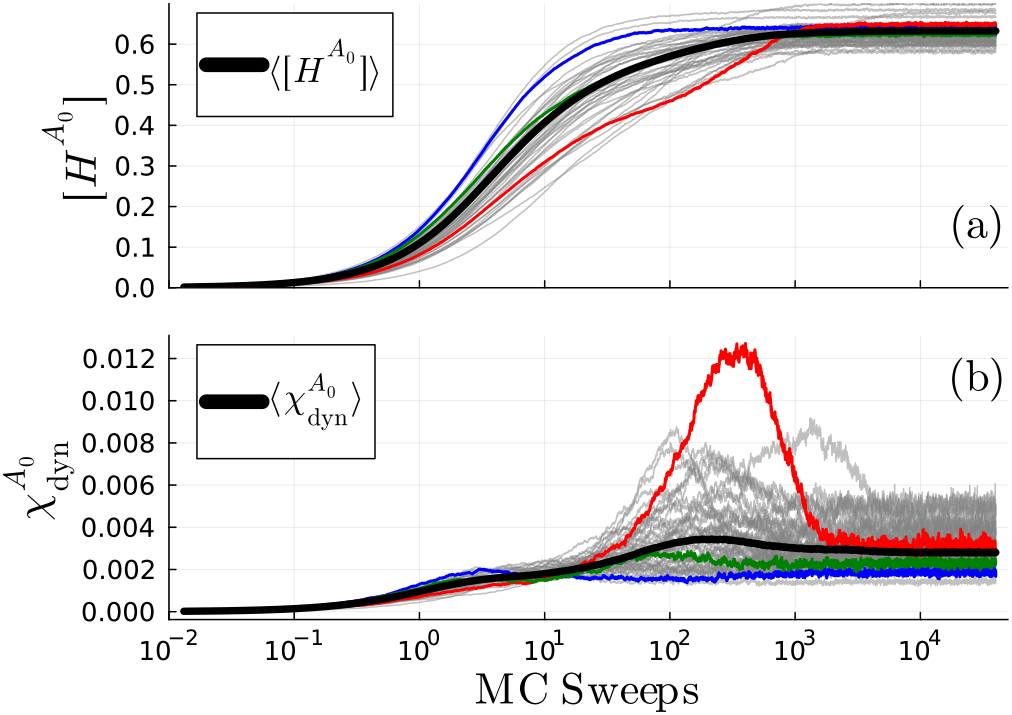
Average (a) and variance (b) of the Hamming distance between the evolving sequence and the ancestor estimated using 10^3^ independent trajectories, for many different ancestors (grey lines). The thick black line represents the average over ancestors, i.e. 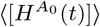 and 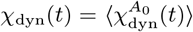, over 200 ancestors. The green, blue and red lines highlights the same specific choices of ancestor used in Fig. 1.

Before describing three interesting cases that we highlighted with colors, we discuss the general traits of such dynamics. The average Hamming distance and its variance are both initially null because all chains start from the same ancestral sequence. Because the DCA model is generative and the evolutionary dynamics respects detailed balance, at large times we expect the simulated sequences to be independent samples from the DCA model, which then reproduce statistical features of the natural ones used for training. This means that the average of 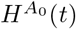 and its variance converge, respectively, to the average and the variance of the Hamming distance between the chosen ancestral sequence and the rest of the natural ones. These values can vary significantly (the final average Hamming distance in Fig. 2a varies from roughly 0.55 to 0.65) depending on how close *A*_0_ is to the other sequences in the natural MSA. The different curves, corresponding to distinct ancestors *A*_0_, show a wide range of behaviors and time scales: some trajectories reach the steady state rapidly while others take much longer, displaying intermediate plateaus as a hallmark of epistasis. In some cases 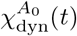 displays a peak and then decreases, while in others the equilibrium value is reached in a monotonic way. The large value reached by 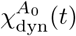 for some initial sequences implies strong fluctuations between distinct evolutionary trajectories, making it hard to predict the dynamics.

In both panels of Fig. 2, we highlighted with colors some representative curves. In particular, the green curve corresponds to an ancestor that behaves in a rather ‘typical’ way, close to the average. The blue curve corresponds to an ancestor that has less epistatically constrained sites, hence the dynamics is faster and less heterogeneous. Finally, the red curve corresponds to a highly epistatically constrained ancestor, which leads to a slower dynamics with an intermediate plateau in 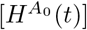, and a large peak in 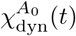. The same three ancestors have been used to construct the PCA plots in Fig. 1 that correspond to time *t* = 50 MC Sweeps.

### B. Dynamical heterogeneity across families

In Fig. 3 we generalize our results to many protein families, chosen to have different ranges of sequence length, MSA depth (number of natural sequences) and equilibration time scales. These families are also interesting because for some of them, experimental data from Deep Mutational Scans and in vitro evolution are available.

**Figure 3.**
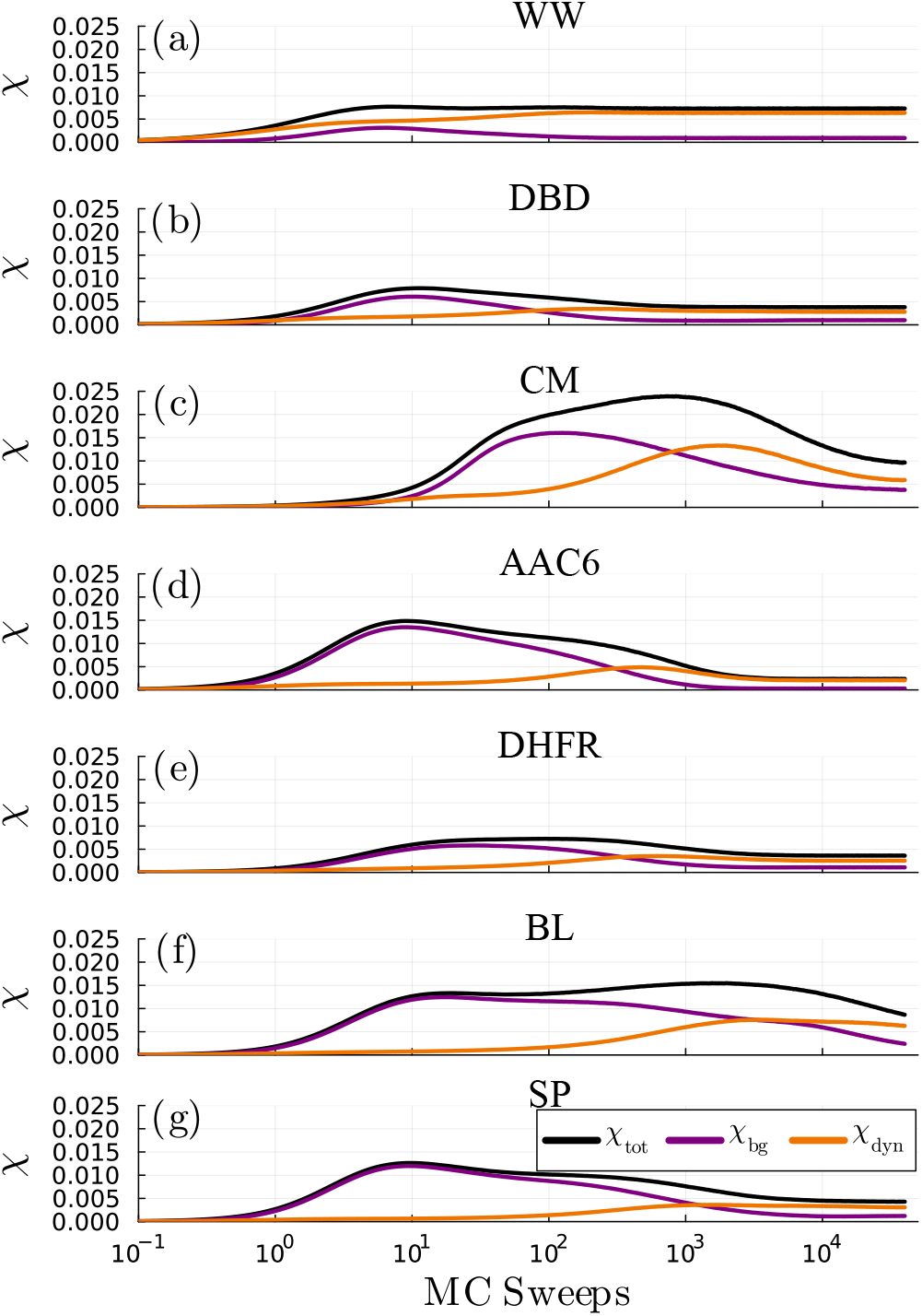
Evolution of the different susceptibilities *χ*_tot_, *χ*_bg_, and *χ*_dyn_, for different protein families. (a) WW domain, (b) DNA-binding domain (DBD), (c) Chorismate Mutase (CM), (d) AAC6, (e) DHFR enzyme, (f) Beta-lactamase (BL), and (g) Serine Protease (SP).

For compactness, we only display the average dynamical fluctuations, with *χ*_tot_ corresponding to the total variance of *H, χ*_dyn_ to the variance due to the stochatisticy of the evolution (averaged over the ancestor), and *χ*_bg_ to the variance due to the choice of ancestor. All the *χ*_tot_ curves display a similar behavior, increasing from zero to a maximum value that is maintained over one or more decades, before finally decreasing to a smaller value corresponding to equilibrium. What matters most to us, however, is the relative importance of the two terms *χ*_dyn_ and *χ*_bg_ in which *χ*_tot_ can be decomposed. The key observation is that background-related fluctuations dominate over dynamical ones at short evolutionary times, while the inverse is true at larger times. This means that, at least over time scales for which *χ*_bg_ ≫ *χ*_dyn_, we can hope to reconstruct with good accuracy the ancestral sequence from which a set of evolutionary trajectories started. Conversely, at larger time scales, the dynamical noise contribution dominates and the trajectory-to-trajectory fluctuations are large enough to hide the signal coming from the ancestral sequence, precluding the possibility to reconstruct it. This behavior is observed in all the protein families that we tested, with the exception of the WW domain in which *χ*_bg_ never grows to larger values than *χ*_dyn_. We attribute this difference to the small size of the WW domain, which prevents the sequences to accumulate enough epistatic interactions. However, both the time scales and the relative importance of the two terms contributing to the total susceptibility vary greatly from family to family (Fig. 3). In DBD, AAC6, DHFR, and SP the peak of *χ*_tot_ is reached quite early in the dynamics and the two contributions *χ*_dyn_ and *χ*_bg_ are almost non-overlapping. On the other hand, for CM and BL, the peak occurs much later due to the two contributions having a significant overlap.

For a given family and a typical ancestor, the time scale at which *χ*_dyn_ and *χ*_bg_ cross, the former becoming larger than the latter, defines a characteristic evolutionary time scale, around which the memory of the ancestor is lost.

### C. Epistatic constraints and dynamical fluctuations

Up to this point, we discussed the time dependence of the different contributions to the total fluctuations. We argued that, as long as the dominant contribution to *χ*_tot_ comes from *χ*_bg_, the ancestral sequence is strongly related to the evolving ones. We now want to better understand the transition to the final regime in which dynamical fluctuations, i.e., *χ*_dyn_, dominate.

In Fig. 4 we show, for the three ancestral sequences highlighted in Fig. 2, how the dynamical part of the susceptibility is strongly related to the epistatically constrained sites (defined as in Sec. III B and in Ref. [29]). To make this relation clear, we first checked that the time scale at which the dynamical susceptibility reaches its peak is compatible with the time scale at which epistatically constrained sites evolve. For a given ancestor *A*_0_, we consider the five most epistatically constrained sites, i.e. those with the largest 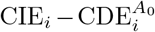. For these sites, we consider at time *t* the frequency of appearance of amino acid 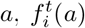, in the set of sequences that evolved from *A*_0_, and from it we compute a time-dependent entropy

**Figure 4.**
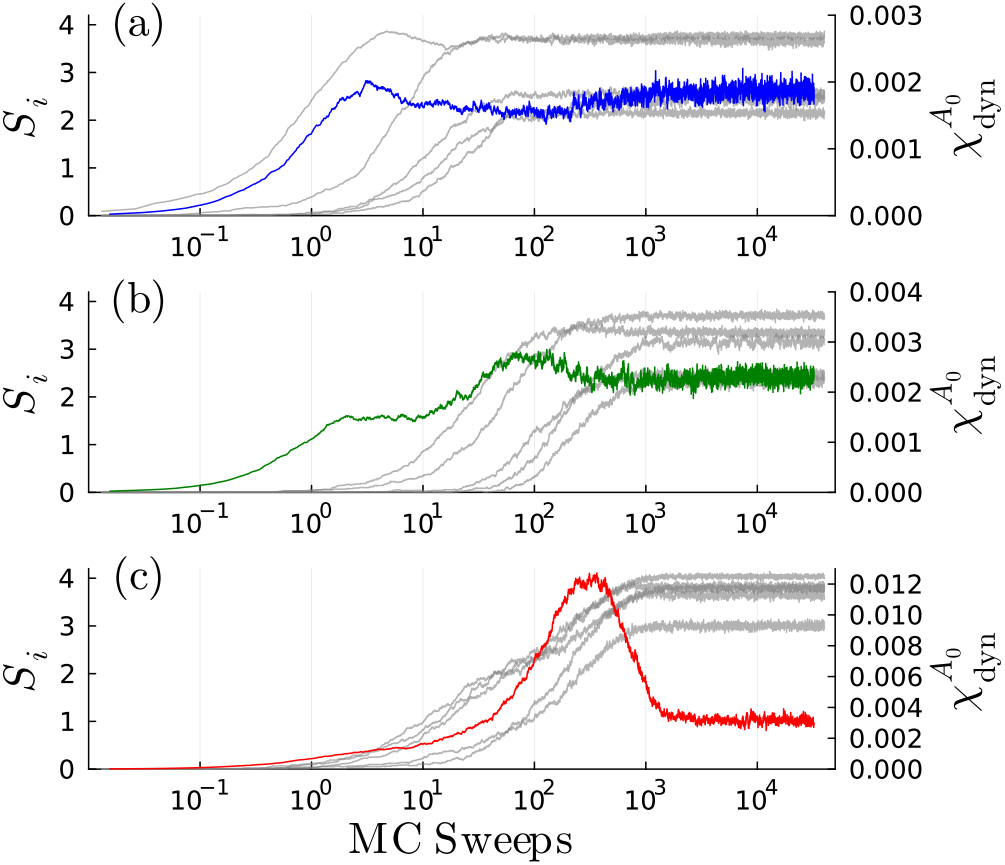
Evolution of the entropy of the most epistatically constrained sites for a specific ancestral sequence of DBD plotted together with 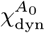 of the same sequence. (a), (b), and (c) are for the blue, green, and red sequences in Fig. 2, respectively.

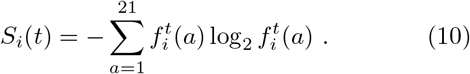

In Fig. 4, 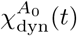 (colored curve) is superposed to *S*_*i*_(*t*) of the five sites (gray curves), as a function of evolutionary time *t*. We observe that the peak of 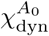 is reached just before the equilibration of the epistatically constrained sites, i.e. the time at which *S*_*i*_(*t*) approaches the CIE_*i*_. The time of equilibration of epistatically constrained sites also increases from Fig. 4a to Fig. 4c, similarly to what happens for the equilibration of the Hamming distance in the same three sequences in Fig. 2. Epistasis does not only affect the time scale at which the peak of 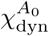 is reached, but also the intensity of the peak, which means that more epistatic ancestors lead to a more heterogeneous dynamics at intermediate times, when the epistatic sites mutate.

To quantify these observations, in Fig. 5 we consider the same set of families as in Fig. 3, namely WW, DBD, CM, AAC6, DHFR, BL, and SP, spanning a wide range of sequence length (from *L* = 31 in WW to *L* = 220 in SP). For each family, we consider a set of ancestors *A*_0_ and we report a scatter plot of the following quantities, each *A*_0_ being a point:

- the value of 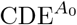 defined as in Eq. (4),
- the maximum value 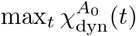, indicated as 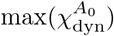 for simplicity,
- and the time *t*_90_ at which the average 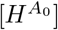 over the trajectories reaches 90% of the equilibrium (*t* → ∞) value.

**Figure 5.**
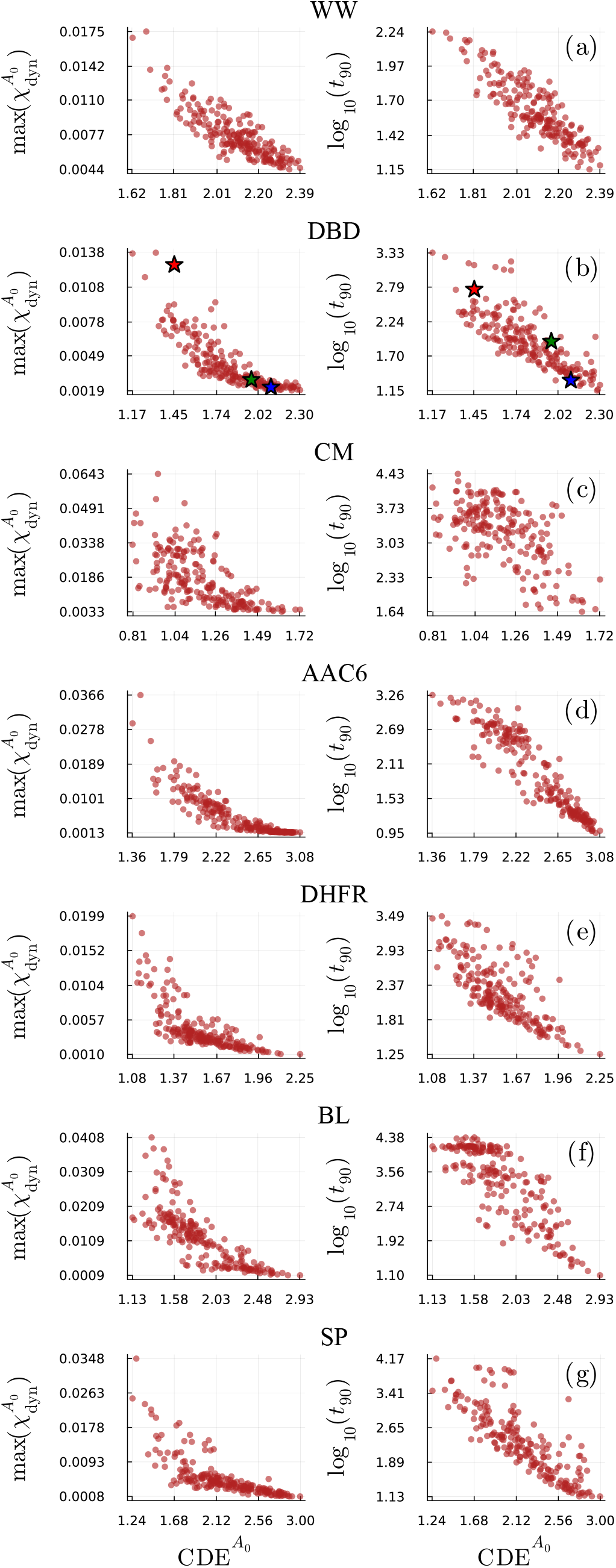
Scatter plot of 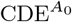 against the maximum of 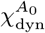 (left) and against log_10_(*t*_90_) (right) for each protein family we considered. For each family, the points correspond to 200 sequences *A*_0_ extracted from the natural MSA with weights (see SM). In (b) colored stars are used to highlight the sequences that correspond to the blue, green, and red curves in Fig. 2.

We observe that both 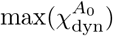 and *t*_90_ markedly increase as 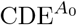 decreases. Hence, a ‘highly epistatic’ sequence with a small value of 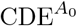 (with respect to the typical value of the family) has many sites with low context-dependent entropy, which cannot mutate at the beginning of evolution, leading to a slower and more heterogeneous overall dynamics. This is for example the case of the red sequence in Fig. 2, for which we have 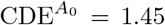, while for the blue one 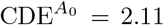. The three sequences considered in Fig. 2 are indicated with stars of the corresponding colors in Fig. 5b.

Epistatically constrained sites also carry the specific signature of the ancestral sequence, because the conserved sites are roughly the same for every sequence of the family and the variable ones carry little to no information. These results thus suggest that tracing back an evolutionary trajectory to its ancestral sequence becomes more difficult as one approaches the peak of *χ*_dyn_, because this is when the epistatic sites, i.e. the sites that carry information about that initial sequence, start to mutate.

### D. Cooperative mutational dynamics

We have established that, when epistatic sites start to mutate, 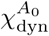 reaches its peak, and that the intensity of this peak is correlated with the amount of epistatically correlated sites in the ancestral sequence. We now want to show that the peak of 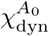 is due to dynamical correlations between different sites, caused by the strong interaction between those sites and the context. These correlations result in a cooperative mutational process, in which epistatically constrained sites can only mutate because other such sites mutate, leading to an avalanche of mutations.

To precisely quantify this effect, one can interpret the dynamical susceptibility as the sum of a site-site dynamical correlation function [47]. The Hamming distance is defined in Eq. (5). Inserting its expression into Eq. (6), we obtain

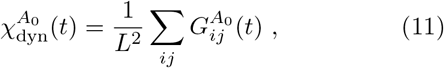

with

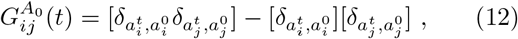

recalling that the Kronecker delta symbol 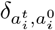 is zero if site *i* is mutated between the two sequences *A*_*t*_ and *A*_0_, and one otherwise. The matrix 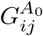 is a time-dependent correlation matrix between sites *i* and *j*. It is high when the sites are dynamically correlated, i.e. if 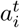 is different from 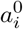, then 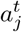 is likely different from 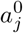 as well, and vice versa. It is low when the sites mutate independently. Because 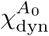 is the sum of 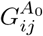 over all pairs (*i, j*), a large value of 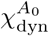 implies that many (*i, j*) are strongly correlated. Furthermore, by looking at this correlation matrix one can infer which pairs of sites mutate in a correlated way during evolution.

The value of the off-diagonal part of 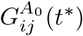, computed at the time *t*^*∗*^ at which a peak in the dynamical susceptibility is observed, is shown in Fig. 6 for six different starting sequences in the DBD family.

**Figure 6.**
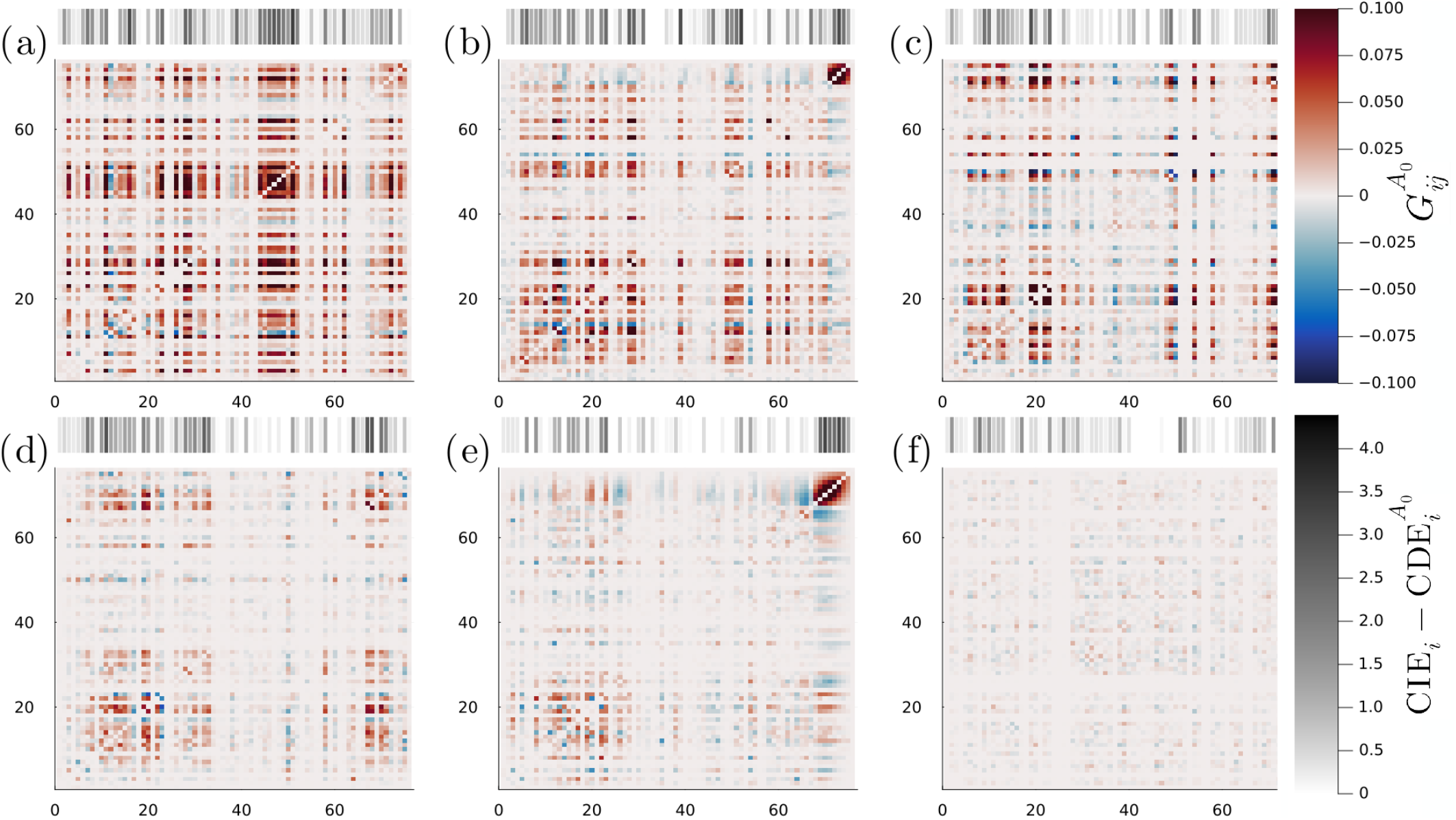
Correlations between sites at time *t*^*∗*^ at which 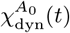 reaches its maximum (with a cutoff at *t*_th_ = 1000 sweeps), for different ancestral sequences. The bar above each snapshot encodes the strength of epistatic constraints for the sites in the ancestral sequence.

Fig. 6a corresponds to an ancestor (red in Fig. 2) for which a large peak and a longer time to equilibrate are observed in the evolution of 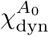. The large value of 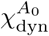, which is just the sum of all matrix elements, implies a large number of positively correlated sites. Fig. 6e corresponds to an ancestor (green in Fig. 2) for which 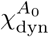 has a smaller peak before saturating. We notice in this case that 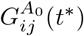 still displays some strong correlations, but the majority of the sites are uncorrelated. Finally, Fig. 6f corresponds to an ancestor (blue in Fig. 2) for which 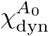 saturates to a small value. In this case correlations are almost absent at *t*^*∗*^, which means that essentially all sites mutate independently. It is thus evident how different initial sequences give rise to different patterns in the 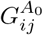 matrix. At large times the chains reach equilibrium and are almost statistically indistinguishable from natural sequences. This means that 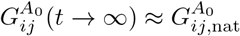, with

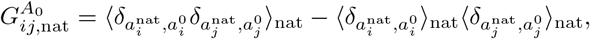

where ⟨… ⟩_nat_ is the average computed over the natural sequences.

In section IV C, we showed that the presence of a peak in 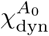 is correlated to 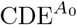, i.e. to the amount of epistatically constrained sites in the ancestral sequence. In this section, we show that these sites also display large dynamical correlations between themselves. The bar above each plot in Fig. 6 is shaded to represent 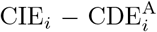, capturing the information about site *i* encoded in the background sequence *A*_\*i*_. This coloring reflects the degree to which each site is constrained by epistatic interactions. The conserved and variable sites are indicated in white, while the epistatically constrained ones are shown in black. As expected, we see good agreement between these residues and the ones that give a large contribution to 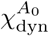. Hence, we conclude that epistatically constrained sites mutate cooperatively around the time scale corresponding to the peak of 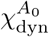.

### E. Response to environmental variation

In the literature on disordered physical systems, it has been established that the dynamical correlations discussed in sections IV C and IV D are related to the linear response of the dynamics to a change in temperature through a kind of fluctuation-dissipation relation [44].

In the present context, the ‘temperature’ is a parameter that controls the probability with which mutations are accepted. High temperature corresponds to all mutations being accepted (very low selection), while low temperature corresponds to only the most beneficial mutations being accepted (very strong selection, i.e. directed evolution). The value of *T* = 1 corresponds to the conditions at which the DCA model is trained on natural sequences, hence *T* ~ 1 corresponds to a neutral drift dynamics during which the neutral space of sequences that have comparable fitness to natural ones is explored [25]. Indeed, Ref. [25] has shown that two distinct in vitro evolution experiments, realized with selection pressures comparable to the natural one, can be described by fitting the temperature in a range slightly above *T* ~ 1.

Hence, in our modeling framework, a change of temperature corresponds to a change of selection strength. Following Ref. [44], we check whether the dynamical susceptibilities are related to the response of the average dynamics with respect to the temperature variation. To estimate such linear response, we consider two evolutionary dynamics starting from the same ancestor, one at temperature *T≠* 1 and the other at temperature *T* = 1, and we compare the average number of accepted mutations 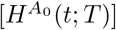 between the two evolutions. The dynamical linear response is given by

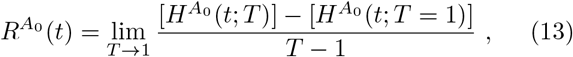

and we emphasize that this quantity depends on the ancestor *A*_0_ and on time *t*. In Fig. 7, we compare the time dependence of 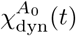 with 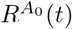 for the same three ancestors as in Fig. 2. Note that we cannot take the limit *T* → 1 due to statistical noise, and we estimate 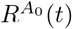 by observing that the curves for several values of *T* close to 1 are almost superimposed to each other. Fig. 7 shows that the fluctuation-dissipation relation seems to hold quite well, such that 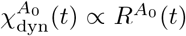, at least for times *t* smaller than the peak of both quantities.

**Figure 7.**
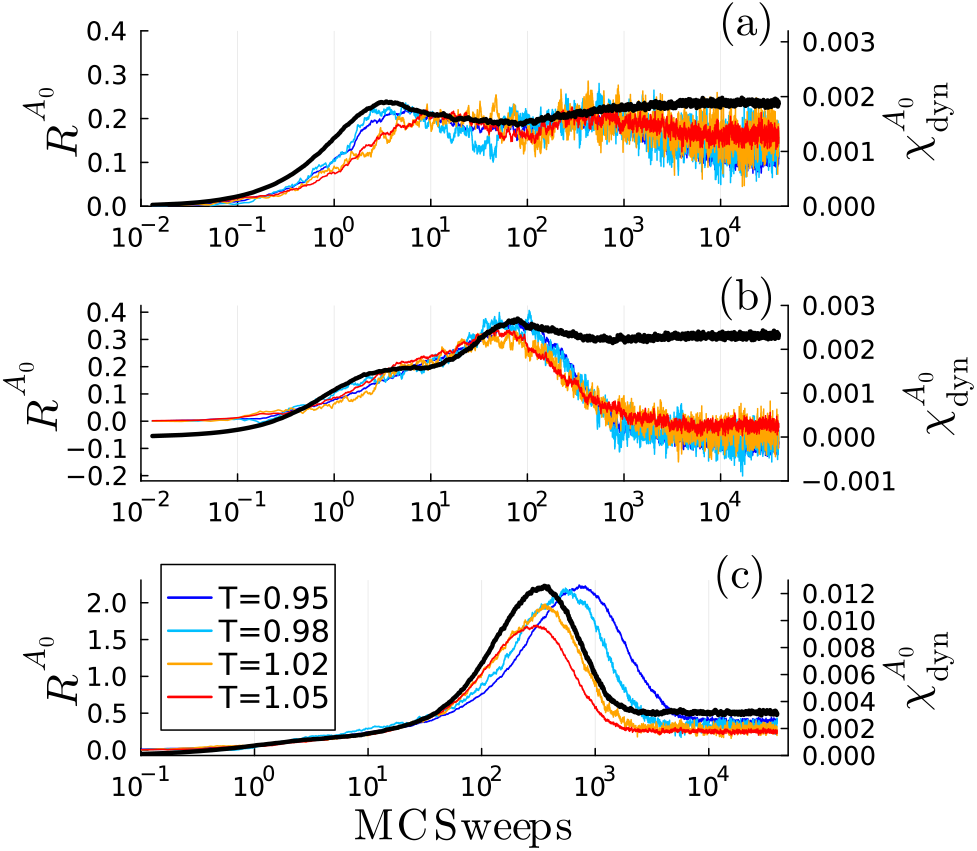
Evolution of the dynamical susceptibility 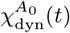 compared with the linear response of the average Hamming distance to a change of temperature 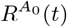. The three panels correspond to the three ancestral sequences highlighted in Fig. 2, (a) blue, (b) green, (c) red.

The results in Fig. 7 suggest that more epistatically constrained ancestral sequences, which display a stronger peak in the dynamical susceptibility, will also display a stronger response of the evolutionary dynamics to a small change in selection strength. Such response is stronger around the time of the peak in 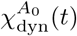, which is also the time at which epistatically constrained sites mutate cooperatively, see section IV D. In other words, *sequences with more epistatically constrained sites are more sensitive to environmental changes*. In the SM, we give an additional analytical argument based on the short-time dynamics, that supports the same conclusion.

### F. Limits of ancestral sequence reconstruction

In an evolutionary process, predictions can be made in two directions: either starting from the ancestral sequence and predicting its future evolution over a given time; or starting from a set of evolved sequences and attempting to infer the ancestral sequence from which the evolution began. An attempt at predicting future evolution has been made in Ref. [25] (see Refs. [51–55] for related studies in disordered systems), where however the prediction was limited to small evolutionary time scales. Because we have repeatedly hinted that our analysis can provide insight into the time scale over which reconstructing the ancestral sequence of an evolutionary process is possible, before concluding, we consider here the ancestral sequence reconstruction problem more explicitly.

For each of the three sequences we analyzed in Fig. 2, we simulate 200 independent evolutionary trajectories, thus obtaining several MSAs of 200 evolved sequences at different evolutionary times *t*. Then, we use the FastML online tool [80] to infer the ancestral sequence starting from those MSAs and using a star-shaped phylogenetic tree. We then check how close the reconstructed sequence is to the actual ancestor.

The results are shown in Fig. 8a, where we plot the evolution of the average Hamming distance for the same sequences as in Fig. 2, and we compare it with the Hamming distance (i.e. the percentage of errors) between the ground truth ancestor and the reconstructed one. At the beginning of the dynamics the FastML tool is able to reconstruct perfectly the original sequence in all of the three cases. This is not surprising, as at short times few mutations appear. The situation is similar at large times, where the MSAs generated from the three starting sequences are statistically indistinguishable and the FastML tool performs equally bad in all of them [81, 82]. Thus, it is more insightful to look at intermediate times, where a significant difference between the evolving sequences is present. After ~10 sweeps the variable sites begin to mutate and the ancestral sequence reconstruction tool cannot reproduce the initial sequence perfectly. The leap in the error percentage with respect to the previous point is however larger for the blue sequence, which is the one with fewer epistastic sites, while the red one is still recovered reasonably well. This difference is present for the subsequent points as well, until the error saturates. In Fig. 8b we show that the Hamming distance (percentage of errors) of the sequence reconstruction results grows roughly proportionally to the average sequence divergence of the evolutionary trajectories for all of the three studied sequences, but more so for the more epistatic red sequence. We conclude that, at least with the FastML tool, the possibility of reconstructing ancestral sequences is mostly determined by the Hamming distance of the ancestor from the evolving sequences, i.e. 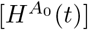, with more errors accumulating when using datasets of more diverged sequences.

**Figure 8.**
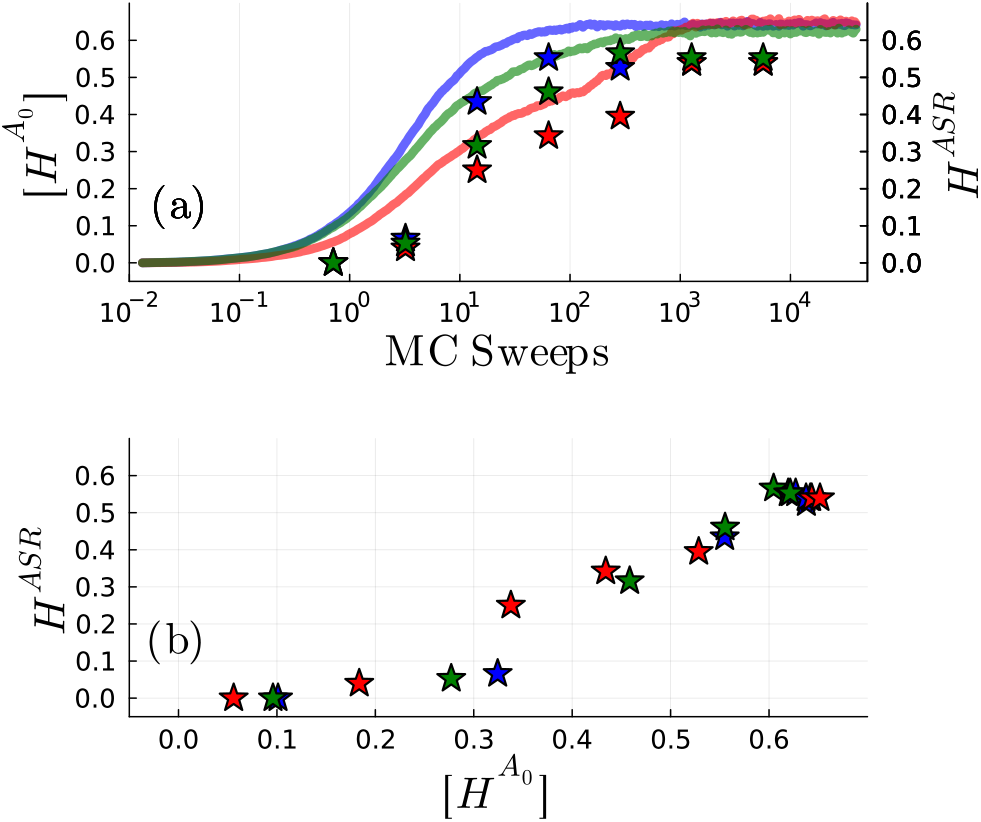
(a) The full lines represent the evolution of the average Hamming distance for the same three ancestors as in Fig. 2. The star symbols represent the Hamming distance *H*^ASR^ between the ground truth ancestor and the one reconstructed using the simulated MSAs at different times. Hamming distance *H*^ASR^ between the reconstructed ancestor and the ground truth as a function of the average distance 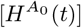 between the ancestor and the MSA used for the reconstruction. The colors refer to the same sequences of panel (a).

Our analysis shows that, for a given evolutionary time, the amount of diversity 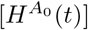 depends strongly on the ancestor, due to highly non-trivial epistatic dynamical correlations. More epistatically constrained ancestors give rise to less diversity, thus allowing for reconstruction over longer evolutionary times (Fig. 8a). Yet, at comparable amount of diversity, more epistatic ancestors are more difficult to reconstruct (Fig. 8b), at least using the FastML algorithm that neglects correlations between sites [83]. These results confirm once more that the evolutionary time limit for predictability is linked to the presence of epistatic sites that carry information about the starting sequence only up to a certain sequence divergence. We further observe that more advanced tools (yet to be fully developed) for ancestral sequence reconstruction might exploit the epistatic correlations to achieve a better reconstruction [84], which is not done in FastML and similar algorithms. We expect that such tools would perform better on more epistatic ancestors such as the red one in Fig. 8.

## V. CONCLUSIONS

In this study, we explore the role of disorder and stochasticity in protein evolution, focusing on the interplay between epistasis, site correlations, and sequence-dependent fluctuations. We simulated evolution in silico using DCA to model the fitness landscape, and an MCMC algorithm to mimic the process of mutation and selection, a methodology that has been validated in previous work [24, 25, 29]. By employing tools from statistical physics [40–49], we quantified how the ancestral amino acid sequence significantly influences early evolutionary dynamics, as measured by the dynamical susceptibility. Our main results are the following.

- Our analysis shows that, during the initial phase of evolution, there is very significant heterogeneity in the dynamics, depending on the choice of ancestral sequence. Some ancestral sequences lead to a smooth evolution where each site mutates almost independently of the others, while others lead to a more complex dynamics characterized by intermediate plateaus and significant fluctuations (Sec. IV A).
- We have shown that, for a variety of protein families, the noise arising from the starting sequence dominates over stochastic evolutionary fluctuations, implying that one can, in principle, trace the evolutionary trajectory back to its origin (Sec. IV B). This effect is more or less pronounced, depending on the family, and our tools allow one to quantify it. However, as time progresses and epistatically constrained sites evolve, this traceability is lost due to the growing influence of the mutational stochasticity.
- The heterogeneity of the initial evolutionary dynamics can be traced back to the amount of epistatic constraints in the ancestral sequence. We introduced a quantity, the average over sites of the context-dependent entropy, and we have shown that this quantity is strongly correlated with the time at which dynamical heterogeneity reaches its peak and with the strength of the fluctuations at the peak (Sec. IV C). This method for assessing sequence evolvability could guide experimentalists in selecting an appropriate starting sequence for experiments, based on the desired functional behavior.
- The amplitude of the global fluctuations can be expressed in terms of a sum of pairwise dynamical correlations between pairs of residues. More epistatically constrained ancestral sequences show groups of residues that mutate collectively, and those patterns can be identified from the correlation matrix 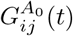 we introduced in Sec. IV D. The observed correlations between sites thus reflect a complex epistatic landscape, where certain residues evolve in a highly context-dependent manner. These correlations could also be leveraged in experimental settings; for instance, one may imagine to control the evolvability of a specific residue by targeting sites that exhibit strong correlations with it.
- We demonstrated that these epistatic sites, which evolve over long time scales, significantly affect the response of the dynamics to a change in environmental conditions. More epistatically constrained sequences lead to a larger response to a change in environment, suggesting potential implications on sequence evolvability imposed by environmental pressures, such as antibiotic concentration (Sec. IV E).
- Finally, we presented a preliminary study of the performance of an Ancestral Sequence Reconstruction (ASR) algorithm (here, FastML), in light of our previous findings. We found that more epistatically constrained ancestors lead to less diversity at comparable time scales, which facilitates their reconstruction. Yet, at comparable diversity, they are more difficult to reconstruct. We believe that this analysis will be instrumental in improving the efficiency of ASR algorithms, which could in principle exploit the correlations identified in this work.

More generally, our findings extend the analogy between protein evolution and disordered physical systems, reinforcing the idea that protein dynamics exhibit characteristics of complex and strongly correlated systems. However, when comparing these results with statistical physics models exhibiting glassy dynamics, we also found qualitative differences, emphasizing the unique constraints imposed by natural evolution on proteins. Future work should aim to further elucidate the connection between epistasis, evolvability, and environmental selection, which may offer insights into evolutionary strategies across diverse biological systems. Moreover, more realistic evolutionary dynamics could be considered. In this paper we focused on independent Monte Carlo chains, which corresponds to evolution on a star tree. Introducing a more complex tree structure could affect the results presented here, as the varying distance between the leaves of the tree and its root and the correlations coming from a common ancestor may affect the computation of 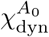. Furthermore, we believe that most of the ideas presented in this work will soon be amenable to experimental testing, thanks to the increased power of in vitro evolution platforms.

## ACKNOWLEDGMENTS

We thank Maria Chiara Angelini, Ludovic Berthier, Pierre Barrat-Charlaix, Simona Cocco, Dongkyu Lee, Arvind Murugan, Clément Nizak, Misaki Ozawa, Andrea Pagnani, Olivier Rivoire, Joe Thornton, Nobuhiko Tokuriki, and Alya Zeinaty for fruitful discussions. This research has been supported by first FIS (Italian Science Fund) 2021 funding scheme (FIS783 - SMaC - Statistical Mechanics and Complexity) from MUR, Italian Ministry of University and Research and from the PRIN funding scheme (2022LMHTET - Complexity, disorder and fluctuations: spin glass physics and beyond) from MUR, Italian Ministry of University and Research.

## SUPPLEMENTAL MATERIAL

### S1. CONSTRUCTION OF THE NATURAL MSAS

The alignments of natural sequences used in this work have been constructed as follows. We believe that the details of the alignment procedure do not affect the results presented in the paper.

- WW: we downloaded the alignment corresponding to the PF00397 family from Pfam [85] and excluded sequences with more than 20% gaps.
- DBD: we used as seed the alignment from Ref. [22] (221 sequences) and ran hmmsearch on uniref90, excluding sequences with more than 20% gaps.
- Chorismate Mutase (CM): we used the alignment from Ref. [79].
- AAC6: we used the same alignment as in Ref. [25].
- DHFR: we downloaded the alignment corresponding to the PF00186 family from Pfam [85] and excluded sequences with more than 20% gaps.
- BL: we used the same alignment as in Ref. [25].
- SP: we downloaded the alignment corresponding to the PF00089 family from Pfam [85] and excluded sequences with more than 20% gaps.

To compute the empirical frequencies used to train the bmDCA model, each sequence is assigned a weight, given by the inverse of the number of other sequences at distance smaller than 20%. The effective number of sequences is the sum of such weights. See e.g. Ref. [78] for details.

In table S1 we show for each family the length of the aligned protein sequences *L*, the depth (number of aligned sequences) *M* of the alignment, and its effective depth *M*_eff_, which serves as a proxy for estimating the diversity of sequences within the alignment.

**Table S1.** Schematic summary of the characteristics of the multiple sequence alignments used for our analysis.

| Family | $L$ | $M$ | $M_{\text{eff}}$ |
| --- | --- | --- | --- |
| WW | 31 | 157809 | 10791 |
| DBD | 76 | 24944 | 3129 |
| CM | 96 | 1259 | 937 |
| AAC6 | 117 | 43576 | 21873 |
| DHFR | 160 | 36612 | 8785 |
| BL | 202 | 18334 | 6875 |
| SP | 220 | 53422 | 22892 |

The three DBD sequences that correspond to the blue, green, and red colors are identified, respectively, with the code

`UniRef90_E4×5B7/28-107`, `UniRef90_O61854/33-105`, and `UniRef90_A0A818T1L9/14-89`. They come, respectively, from the organisms *Oikopleura dioica (Tunicate), Caenorhabditis elegans*, and *Rotaria sp. Silwood1*.

**Figure S1.**
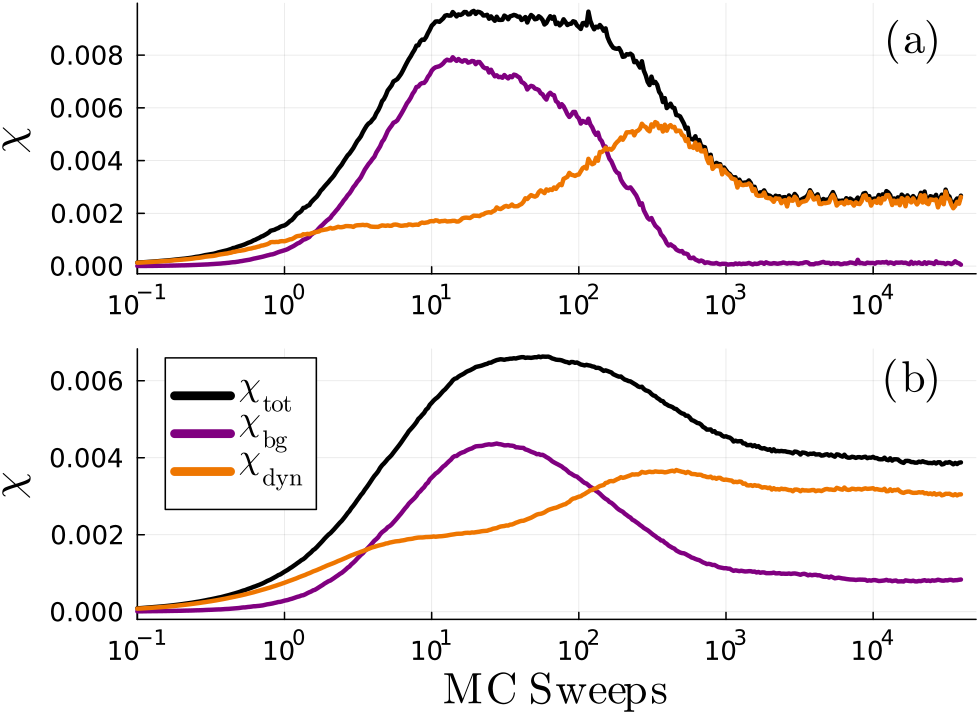
Evolution of *χ*_bg_, *χ*_dyn_ and *χ*_tot_ as a function of Monte Carlo sweeps using a sparse edge-activated version [86] of the Potts model (a) and using the dynamics of Ref. [29] implementing insertions (b), deletions and single nucleotide substitutions for the DBD family.

### S2. DIFFERENT MODELS AND EVOLUTIONARY SCHEMES

Different variants of the bmDCA model that we employed in the main text have recently been suggested. We performed simulations using a sparse edge-activated [86] version for the DBD family, without any significant change in the results (Fig. S1a).

As we already pointed out, the evolutionary protocol that we implemented is a simple Metropolis Monte Carlo defined over amino acid sequences, which is schematically described as follows:

1. Initialize with a protein sequence *A*^0^.
2. At each iteration:

(a)Select a random site *i* ∈ {1, …, *L*}.

(b)Propose a mutation 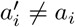 by selecting a new amino acid at site *i* uniformly from the alphabet.

(c)Compute the energy difference:

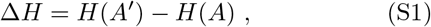

where *A*^*′*^ is the sequence with the proposed mutation.

(d)Accept the mutation with probability

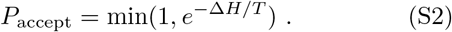

If accepted, update 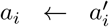; otherwise, keep *a*_*i*_.

3. Repeat for a sufficiently large number of iterations to reach equilibrium.

After equilibration, sequences sampled from this procedure approximate the equilibrium distribution of the Potts model.

This dynamics is far from being realistic, especially at short time-scales. As a matter of fact, a more detailed approach was recently described in [29], where it was shown that an algorithm working with single-nucleotide substitutions (instead of amino acid substitutions), insertions and deletions was able to correctly reproduce the dynamics both at long and short time scales. We checked that this more refined algorithm, which was shown to overperform a simple Metropolis at short time scales [25], produces qualitatively similar results for the quantities of interest for our work (Fig. S1b).

### S3. PROFILE MODELS

As epistatic sites play an important role in our description, it is interesting to study a dynamics in which they are totally absent. This can be done by using a profile model in which only a local field acts on each site, eliminating the coupling between different protein residues. Such profile models can be built in two ways, analogously to what has been done in Ref. [29]. We call *global profile* the model in which the local field is set equal to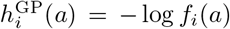, in such a way as to reproduce the one-point frequencies. In this case, the sites evolve following their local field independently from one another. In Fig. S2a we show how the dynamical susceptibility evolves for the same three sequences we studied in the main text. We saw above that the blue sequence had a smaller 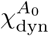 and that it converged to the long-time limit quite fast. Differently, the red sequence showed a large peak and only converged after many sweeps. In this figure we see that when using a global profile model the starting sequence does not affect the 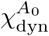 anymore. As the interactions are set to zero, the initial condition plays a much less important role and all the curves seem to behave in a rather similar way, showing a small peak at a homogeneous time scale. Couplings between residues and hence epistasis are needed to observe the qualitative behavior presented in the main text.

**Figure S2.**
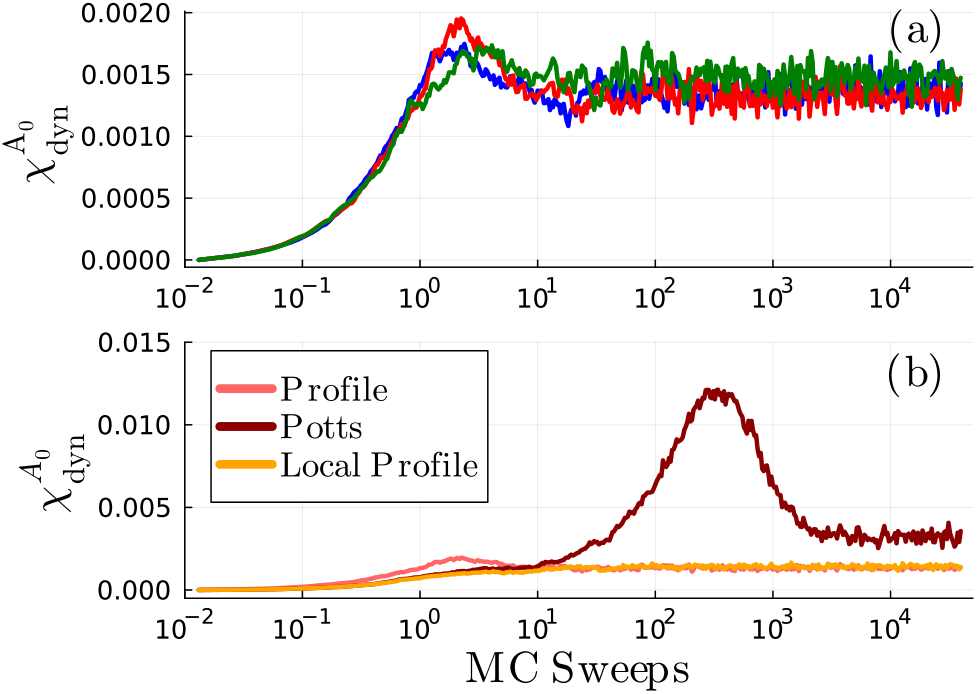
Evolution of the dynamical susceptibility for the profile models. In (a) we show the evolution of 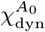 for the same starting sequences used in the main text studied with the global profile model, while in (b) we compare the results for the red sequence obtained using the two profile models and the full one.

One can also define a *local profile* model, in which the fields acting on the residues are obtained as

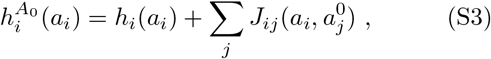

and therefore depend on the initial sequence *A*_0_. We compare both these profile models and the Potts one Fig. S2b for one sequence *A*_0_. At the very beginning of the dynamics the three models are very similar, as only few mutations occurred. Already after few steps, less than a sweep, the global profile model displays the small peak discussed above, due to the uncorrelated mutations occurring at different residues. The local profile takes into account the role of the initial condition and hence it follows a dynamics similar to the full Potts model for a longer time (~10 sweeps). However, for even longer times, in the full Potts model the evolution of the background changes the local field acting on each residue, allowing sites to evolve in a coordinated manner. This results in the presence of the peak, as discussed in the main text. In the local profile this correlated evolution cannot take place, as the local fields are kept constant. Hence, the dynamical susceptibility quickly saturates to its long time limit value.

### S4. DIFFERENT DEFINITION OF SUSCEPTIBILITY

It is also interesting to consider a different definition of susceptibility. In particular we already defined 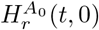 as the Hamming distance between the sequence that evolved for *t* steps under the predefined evolutionary noise *r* starting from the initial sequence *A*_0_ and the initial sequence itself. We can define two supplementary Hamming distances, this time computed at fixed Monte Carlo sweeps between different independent simulations of the system. In particular we define

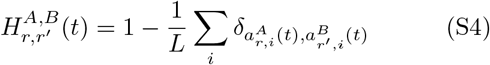

as the Hamming distance between sequences that have independently evolved from different initial sequences *A* and *B* for the same time *t*. The […] average is then computed by summing over all different realizations of the couples of evolutionary noises (*r, r*^*′*^), while the ⟨… ⟩ average is computed by summing over all couples of different initial sequences (*A, B*). It is interesting to compare the particular case 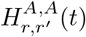 with what we showed before. Because our dynamics is time-reversible, we expect that 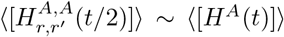. We checked that this is indeed the case.

In the main text we showed that 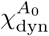 can be expressed as the sum of a correlation function between sites (see Eqs. 11 and 12). If two sites both mutated after a time *t* with respect to the initial sequence *A*_0_, then they presented a high correlation. In this case, 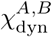 can be still seen as the sum of site correlations, but, differently from the previous case, two sites *i* and *j* are highly correlated if after time *t* the realizations starting from the two sequences *A* and *B* display the same amino acids in those sites, namely

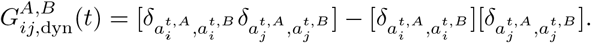

Analogously to what is presented in the main text, we can define 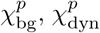 and 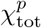 where *p* apex indicates that we are considering a pairwise metric. Because 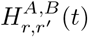 strongly depends on which couples *A, B* are chosen to initialize the two dynamical evolutions, we can easily expect that the impact of the background initialization 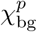 will be dominant at the beginning of evolution. Instead, when equilibrium is reached, 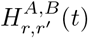 tends to the mean pairwise hamming between the sequences in the training set as the model is generative. Hence, at later times the memory of the initial condition is lost and 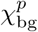 vanishes. This is confirmed by Fig. S3.

**Figure S3.**
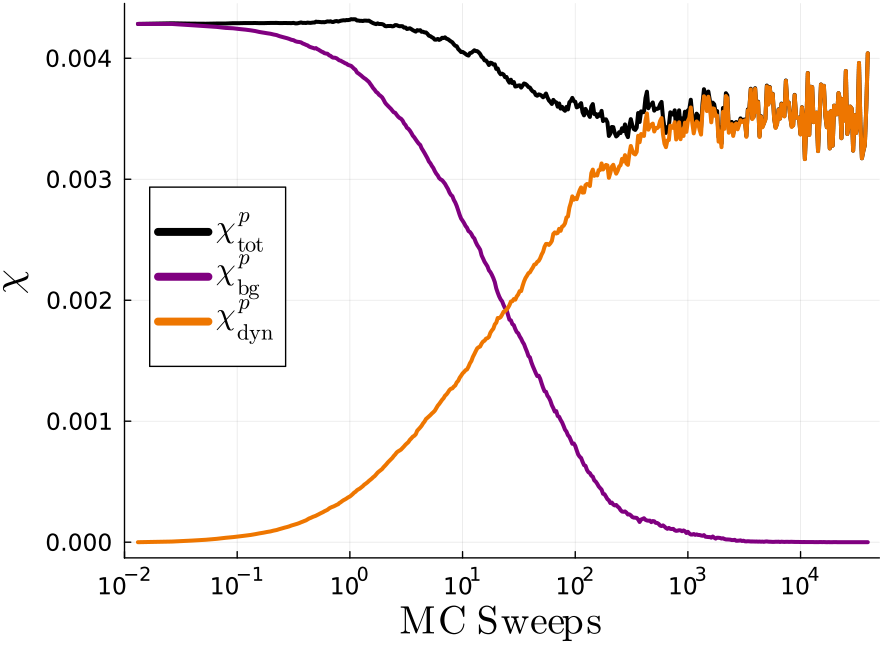
Evolution of 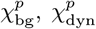 and 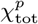 as a function of Monte Carlo sweeps for the DBD family.

### S5. RESPONSE TO ENVIRONMENTAL VARIATION

To complement the analysis presented in the main text concerning the response to a change in selection pressure, we checked how much the mutability of the ancestral sequence 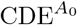 changes with respect to a shift in selection pressure. In Fig. S4 we plot such derivative as a function of 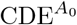 for a set of sequences of the DBD family. It is clear that epistatically constrained sequences have a higher change in mutability when subject to a change in selection pressure (i.e. a change in the simulation temperature). As a matter of fact, the average context-dependent entropy of a given sequence *A*_0_ can be expressed as a function of the local fields 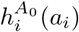 defined in Eq. (S3) as follows

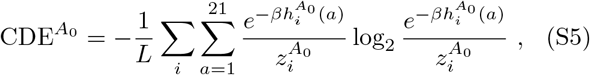

with 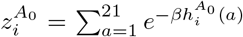. Hence, the derivative with respect to inverse selective pressure can be analytically computed as follows:

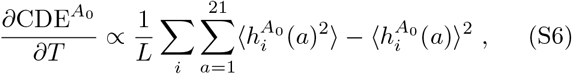

where 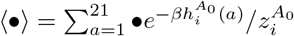. This expression indicates that sites having a greater variance of local fields are more prone to change their mutability as a sudden change in selective pressure takes place. This analysis offers interesting insights into the evolvability of protein sequences which should be corroborated by evolution experiments.

**Figure S4.**
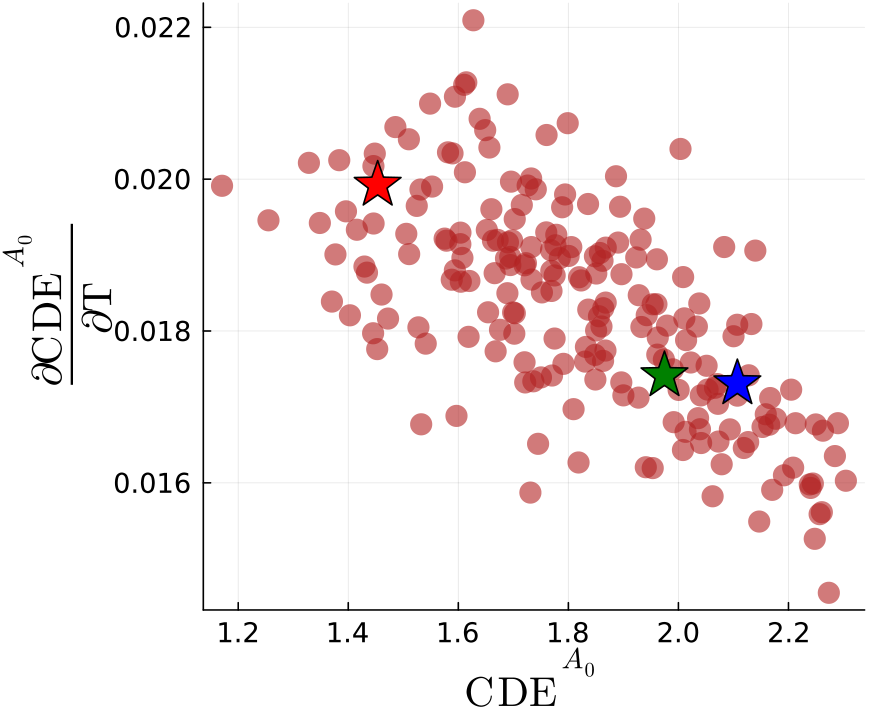
Derivative of the context-dependent entropy with respect to temperature (i.e. inverse selection pressure), as a function of the sequence mutability 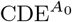 for 200 sequences of the DBD family. The colored stars refer to the three sequences of Fig.2.

